# BERTMHC: Improves MHC-peptide class II interaction prediction with transformer and multiple instance learning

**DOI:** 10.1101/2020.11.24.396101

**Authors:** Jun Cheng, Kaïdre Bendjama, Karola Rittner, Brandon Malone

## Abstract

**Motivation:** Increasingly comprehensive characterisation of cancer associated genetic alteration has paved the way for the development of highly specific therapeutic vaccines. Predicting precisely binding and presentation of peptides by MHC alleles is an important step towards such therapies. Recent data suggest that presentation of both class I and II epitopes is critical for the induction of a sustained effective immune response. However, the prediction performance for MHC class II has been limited compared to class I.

**Results:** We present a transformer neural network model which leverages on self-supervised pretraining from a large corpus of protein sequences. We also propose a multiple instance learning (MIL) framework to deconvolve mass spectrometry data where multiple potential MHC alleles may have presented each peptide. We show that pretraining boosted the performance for these tasks. Combining pretraining and the novel MIL approach, our model outperforms state-of-the-art models for both binding and mass spectrometry presentation predictions.

**Availability:** Our model is available at https://github.com/s6juncheng/BERTMHC

## 1 Introduction

Adaptive immunity is the mechanism by which a long lasting, specific protection is acquired by the body after exposure to a pathogen such as a microbial agent or a cancerous cell. Adaptive immunity is triggered by clonal expansion and onset of lymphocytes. Adaptive immunity is classically divided into two components: the humoral immunity and the cellular immunity (Janeway *et al.*, 2001). While the immoral immunity relates to the onset of antibodies from mature B cells, the cellular immunity is the results of the expansion of target specific T-cell population. Priming of T and B cell response against a given antigen is dependent on presentation of the said antigen by HLA class II molecules on the surface of professional antigen presenting cells (Lang *et al.*, 2001). Detection of the MHC class II–Antigen complex by a T-cell will trigger a signalisation cascade, leading to the maturation of T cells in a helper T cell that will stimulate both the onset of an effective T cell response and the maturation of pathogen specific B-cell (Al-Daccak *et al.*, 2004). The onset of helper T cells is critical for development of a sustained adaptive response. As such, class II presentation is critical in vaccine development when attempting to leverage the immune system for therapeutic purpose. To ensure that a large repertoire of diverse pathogen antigens are presented, class II MHC are encoded by genes characterized by a very high level of polymorphisms (Neefjes *et al.*, 2011). Furthermore, each individual has several alleles over several loci encoding for class II MHC molecules. These polymorphisms lead to the presentation of different specificity and different presented repertoire across individuals.

This high level of polymorphism is an evolutionary trait allowing the immune system to react against a very large number of pathogens at the individual level. The corollary implication of this high level of polymorphism is that each individual will eventually present different peptides. This has critical implications when attempting to design vaccines and select antigens. Historically, vaccine development involved the use of large antigens, representative of the target pathogen. Because of their size, these antigens are likely to contains peptides sequences that will be presented by a wide spectrum of HLA genotypes. More recent vaccines, and in particular vaccines directed to cancer cells, are targeted at restricted sequences as the pathogen - the cancer cell - is minimally different from the host cell (Tanyi *et al.*, 2018). In order to increase vaccine efficiency, it is critical to identify sequences that are likely to be presented by the class II HLA in the individuals that will receive the vaccine.

The presentation of peptides to T cells involves a series of processes. Important steps include binding between MHC molecules and peptides, as well as presentation of the MHC–peptide complex to the cell surface. Experimental assays have been developed to study and quantify many of these processes. The binding affinities between MHC molecules and peptides can be measured by *in vitro* binding assays (Sidney *et al.*, 2013). Mass spectrometry can be used to detect peptides eluted from the cell surface to determine peptide presentation (Purcell *et al.*, 2019). Thousands of data points have been generated by such assays for hundreds of different MHC molecules (Vita *et al.*, 2019). However all these wet lab methods remain labour intensive and subject to a number of biases.

Given the importance of the problem and the availability of the data, many methods have been developed to predict MHC–peptide binding and peptide presentation (Peters *et al.*, 2020). In some approaches, a single model is trained specifically for each MHC allele; other approaches instead train a single model (a *pan model*) covering all MHC alleles. The prediction performances of MHC class I models have reached a high level (auROC *>* 0.95, Peters *et al.*, 2020; O’Donnell *et al.*, 2020; Reynisson *et al.*, 2020b). On the other hand, models for class II still have limited performance. Despite recent progresses (Reynisson *et al.*, 2020b), there is still a need for better performing models. One significant limiting factor for MHC class II models is the limited amount of training data compared to class I. Thus, models that can efficiently use all of the limited available data and transfer knowledge from other sources are extremely valuable.

Recent advances in natural language processing have enabled techniques to train complex models that understand semantics from text without labels (self-supervised learning) (Peters *et al.*, 2018; Devlin *et al.*, 2019). Such models are trained to predict words masked out in a sentence or to predict the next word or sentence following some context. Similar techniques have also been applied to proteins (Rao *et al.*, 2019; Nambiar *et al.*, 2020; Heinzinger *et al.*, 2019). Since these models do not require labels to train, they can be trained on very large corpora of protein sequences across many species. One example of these models is the TAPE model (Rao *et al.*, 2019), which was trained with 31 million protein sequences from the Pfam database (El-Gebali *et al.*, 2019). The model has been shown to be helpful in a variety of downstream tasks such as remote protein homology prediction (Fox *et al.*, 2013) and stability prediction (Rocklin *et al.*, 2017). Detailed analysis of the model has shown that it captures long-range interactions in 3D structure (Vig *et al.*, 2020). It is highly relevant to explore whether pretrained protein sequence models can be helpful for MHC–peptide binding and presentation prediction, especially for MHC class II where much less data is available.

As mentioned, each person has multiple MHC class II molecules; thus, typical mass spectrometry experiments cannot precisely identify the MHC molecule to which a peptide was bound. In designing personalized vaccines, we are also interested in the likelihood of response given all the MHC alleles an individual carries. Therefore, it is important to develop algorithms to predict the likelihood of presentation for a peptide given a set of MHC molecules.

Here, we focus on developing models for predicting MHC–peptide binding and presentation for MHC class II. We show that models taking advantage of self-supervised pretraining from large corpora of protein sequences can achieve better performance on both binding and presentation prediction tasks. We found pretraining to be extremely valuable in the case where training data is limited. Additionally, we propose a novel multiple instance learning algorithm to account for the limitation that mass spectrometry data often cannot precisely identify the exact MHC molecule to which a peptide was bound. We foresee our work to be valuable in T-cell-based immunotherapy and provide new directions for training peptide binding models with limited training data.

## 2 Materials and Methods

### 2.1 MHC class II binding data

To train the MHC class II binding model, we used the data from (Jensen *et al.*, 2018) since it has been designed to minimize the overlap between the training and evaluation sets. The original data was collected from the Immune Epitope Database (IEDB, Vita *et al.*, 2019, accessed on 30 June 2020) up to the year 2016. The data consists of 134 281 data points and covers HLA-DR, HLA-DQ, HLA-DP and H-2 mouse MHC allele. The affinity labels were transformed from IC_50_ to values between 0 and 1 with the formula 1 *−* log(IC_50_)*/* log(50 000).

### 2.2 Independent MHC class II binding data

The data from Jensen *et al.* was collected from IEDB up to the year 2016. To benchmark on an independent dataset where no model has been used for training or validation, we collected quantitative binding data from IEDB and filtered out data already used in Jensen *et al*. In addition, we collected additional independent binding data from the Dana-Farber repository (Zhang *et al.*, 2011). In the end, we collected 2 413 additional MHC–peptide pairs covering 47 MHC class II alleles. The complete list of benchmark data is available in Supplementary Table 2.

### 2.3 MHC class II presentation data

To train the MHC class II mass spectrometry presentation model, we used the data curated from Reynisson *et al.* (2020b). The original data were curated from IEDB and other public sources. The data covers 41 MHC class II alleles with peptide lengths ranging from 13 to 21. Each data point consists of the peptide ligand, the source protein and list of possible MHC class II allele bound to the peptide. The data points where only one MHC allele is unambiguously given are referred as *single-allele data* (SA), whereas the data points where multiple potential alleles, are given are referred to as *multi-allele data* (MA). Reynisson *et al.*, selected negative peptides by randomly sampling from the UniProt database. Peptide lengths for the negatives are sampled uniformly from 13 to 21.

### 2.4 Independent MHC class II benchmark presentation data

We also filtered for further independent datasets from IEDB (accessed on 30 June 2020) on which no existing models had been trained or evaluated. All data presented in the training and evaluation set of Reynisson *et al.*, were filtered out. We only kept 8 170 peptides with length between 13 and 21, in line with Reynisson *et al.*, and alleles which have more than 50 positive peptides. To generate negative mass spectrometry decoys, we randomly sampled per allele 10× negative peptides from the human proteome with the same peptide length distribution as the positive peptides. This evaluation data of 95 638 peptides was provided as Supplementary Table 3.

### 2.5 Patient mass spectrometry data

To evaluate our model on an independent set of data, we generated a dataset of HLA class II presented peptides from 6 patients with cancer. Briefly, cancer tissue was collected on surgical material in patients undergoing non-small cell lung cancer resection. The collection of biological material was performed in accordance to international and local clinical research regulation and subject to ethical review boards approvals. Accordingly, participation to the study was contingent to informed consent from individual patients. After tissue lysis, solubilized HLA complexes were purified with antibody-conjugated resin (clone L243). HLA-bound peptides were eluted in acidic condition, further purified and desalted before final mass spectrometric analysis. Eluted preparations were analyzed by data-dependent (DDA) mass spectrometry on an HF-X hybrid quadrupole-Orbitrap mass spectrometer. Peptides were identified with MaxQuant Cox and Mann (2008) (Supplementary Methods). In total 15 277 unique peptides were identified. Between 1 500 and 7 000 unique peptide sequences were identified for the samples (Supplementary Table 1).

### 2.6 Pretrained protein BERT model

We use the pretrained bidirectional encoder representations from transformers (BERT, Devlin *et al.*, 2019) from the TAPE repository to model the input amino acid sequences. The TAPE model was trained with *self-supervised learning* from a dataset of over 31 million protein sequences. Briefly, t aking u nlabelled p rotein s equence as input, the TAPE model was trained with two tasks. One task is bidirectional *next-token prediction* (predicting *p*(*x*_*i*_|*x*_1_, *x*_2_, …, *x*_*i*−1_) and *p*(*x*_*i*_|*x*_*i*+1_, *x*_*i*+2_, …, *x*_*L*_)), and the other task is *masked-token prediction* (predicting *p*(*x*_masked_|*x*_unmasked_)). The model has 12 layers with 12 self-attention heads (Equation 1) in each layer, which enables the model to learn long distance interactions. For an input amino acid sequence ****z**** = (*z*_1_, *z*_2_, …, *z*_*L*_), the output of the model are *L* continuous vectors of dimension 768 corresponding to the input amino acids.

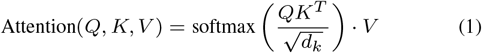

### 2.7 Supervised training of MHC class II models

We trained two BERT models, one each for MHC class II binding and presentation prediction. Both models were initialized with the BERT model parameters from the TAPE repository. Both models are pan models, and they take as input the concatenated amino acid sequences for the MHC pseudosequences (Jensen *et al.*, 2018; Reynisson *et al.*, 2020b) and the peptide (MHC then peptide). Because peptides have variable lengths, we pad all sequences to the same length with an input padding token. The target for the binding prediction model is the (real-valued) binding affinity between the peptide and MHC, while the target for the presentation models is binary (presented or not).

As shown in Figure 1, our model architecture consists of three main components: the BERT component, a pooling layer, and a final multilayer perceptron (MLP) block. The input sequence is first tokenized and used to extract the continuous vectors from the BERT component. The padding token vectors were masked, and the remaining vectors are then pooled by taking the mean over the sequence dimension. This vector is then used as input for the MLP, which consists of two fully connected layers with hidden dimension of 512. The MLP was then used to predict either the MHC binding affinity or cell surface presentation. We used the standard mean squared error loss function for the binding model and the weighted cross-entropy loss functions (weight 10 for the positive class) for the presentation model when training with SA data. The loss function for training MA data is described in Section 2.8. We train the model end-to-end with backpropagation; that is, we also optimize the parameters from the pretrained TAPE model.

**Figure 1.**
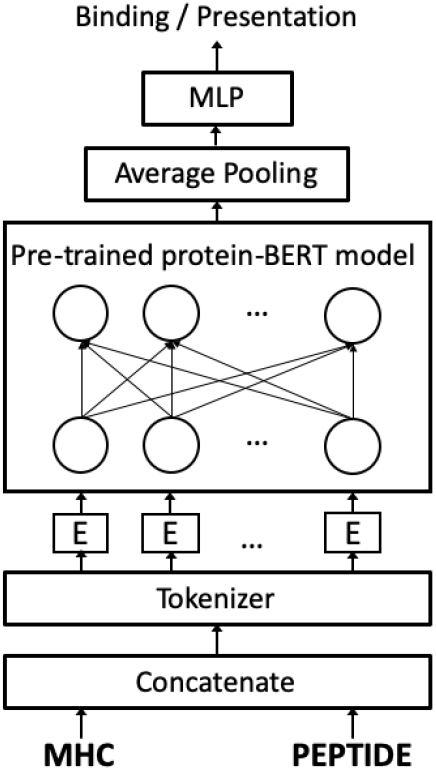
BERTMHC model architecture. Peptides were first encoded into tokens with a tokenizer, where each token is a single amino acid. Each token is then embedded into a 768 dimensional vector. This followed by 12 layers of multi-head self-attention layer with 12 attention heads at each layer. The output of the BERT encoder is then mean-pooled along the sequence dimension. A 2-layer feed forward neural network uses that to predict either binding affinity or presentation.

For both the binding and presentation models, we used random search to search for the best hyperparameter combinations on the first cross-validation fold. We used an initial learning rate of 0.15 for the binding model and 0.01 for the presentation model. All models were trained with stochastic gradient descent with momentum of 0.9; the learning rate was reduced by a factor of 0.1 after 2 epochs of no performance improvement. After identifying high-quality hyperparameters, we use them to train three models with different initializations for each cross-validation fold for the binding model; only one model per fold was trained for the presentation task. The two tasks were trained independently.

### 2.8 Multi-allele deconvolution with multiple instance learning

In the MA presentation data, each peptide is associated with a bag of alleles. The bag is labeled as positive if at least one of the alleles presented the peptide, otherwise the bag is labeled as negative. Training from such data is sometimes referred to as *deconvolution* (Bassani-Sternberg and Gfeller, 2016; Reynisson *et al.*, 2020a,b).

We model training the prediction model *f*_*θ*_ from the MA data as a multiple instance learning (MIL) problem. We denote the *i^th^* bag with *m* alleles as *A*_*i*_ = *{a*_*i*1_, *a*_*i*2_, …, *a*_*im*_} and the corresponding peptide sequence as *s*_*i*_. At each training step, we predict the probability *p*(*y*_*ij*_ = 1|*x*_*ij*_) for every instance (*a*_*ij*_, *s*_*i*_) in the bag as 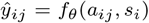 with our neural network model *f*_*θ*_. A symmetric pooling operator **h** is used to calculate the prediction of the bag from the predictions of instances within it.

We further propose to incorporate the confidence of the deconvolution operation. At each training epoch we weight each positive data point *i* from deconvolution by a calibrated predicted probability of being positive 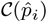. The probabilities are calibrated by performing isotonic regression (Barlow *et al.*, 1972) from the predicted logits and the labels on the training set. We do not weight the negative classes since there is no ambiguity with their labels. Therefore, we compute the loss for the MA data as follow:

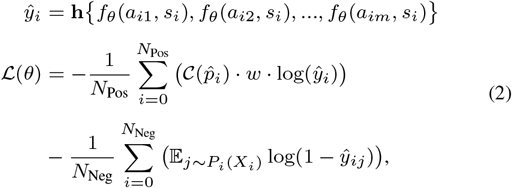

where 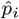is the prediction of *p*(*y*_*i*_ = 1|*A*_*i*_, *s*_*i*_) from the previous epoch of the model, 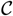is the calibration function, and *w* is the weight for the mass spectrometry positive class for compensating class imbalance in the dataset. We use *w* = 10 for all presentation models. We used max pooling as the pooling operator **h**, although other pooling operations, such as attention-based pooling (Ilse *et al.*, 2018), are also applicable. For computational reason, we perform negative sampling with a probability distribution *P*_*i*_(*X*_*i*_) instead of using all negative samples. We use the most likely positive example predicted by the current model from the negative bag. Therefore the sampling distribution *P*_*i*_(*X*_*i*_) = *P*_*i*_(*x*_*i*1_, *x*_*i*2_, …, *x*_*im*_) is defined as:

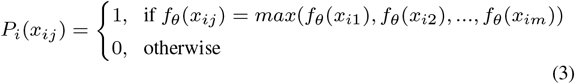

In addition, we follow a two-state training procedure. Before multiple instance training with MA data, we first train the model only with SA data. After 8 epochs, we combine SA and MA data and train them jointly with MIL, where the bag for SA data has only one element. We stop the training process if the model performance (average precision) on the evaluation set do not improve after 5 epochs. Algorithm 1 summarizes the probability reweighted multiple instance learning algorithm.

**Algorithm 1:**
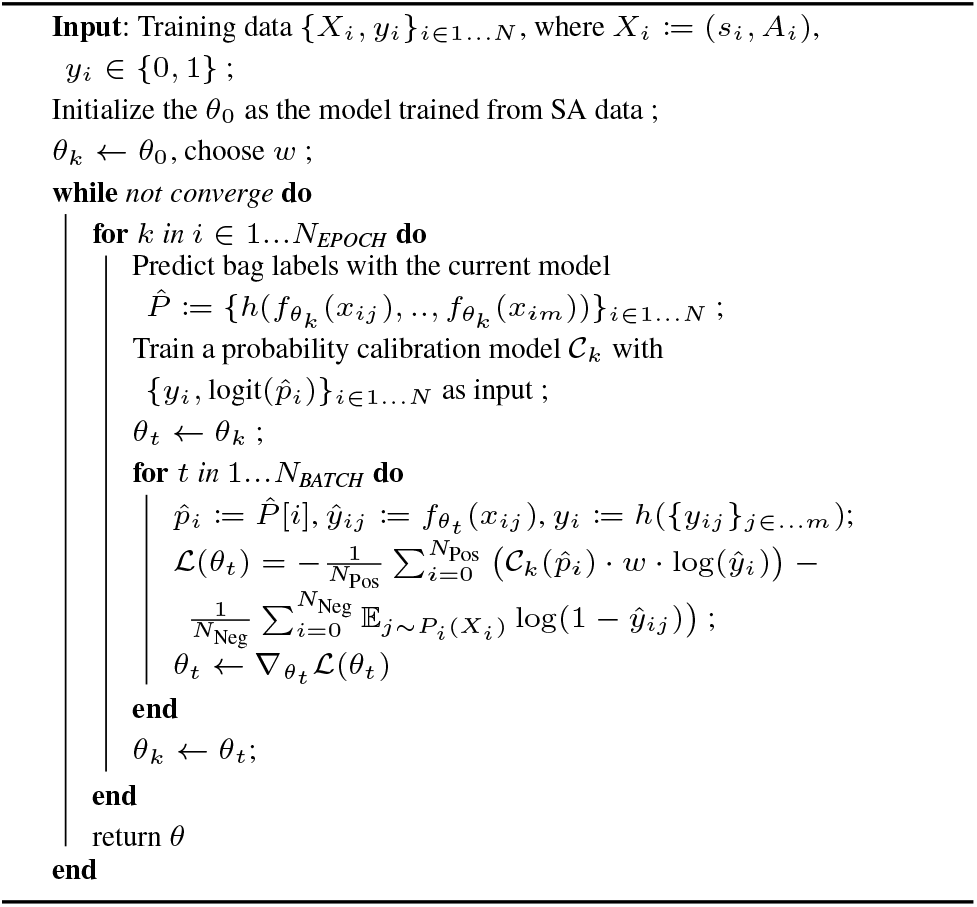
Probability Reweighted Multiple Instance Learning

Predictions on MA data in the test sets were performed as follow: for each bag of alleles and peptide *x*_*i*_ = (*A*_*i*_, *s*_*i*_), predictions 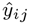 are made with each candidate allele separately. The label of the bag is predicted as

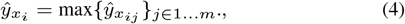

where *m* is the number of potential alleles which may have presented the peptide. 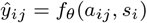 is the model *f*_*θ*_ predicted score for peptide *s*_*i*_ and allele *a*_*ij*_.

Meanwhile, the allele label for the bag is assigned as

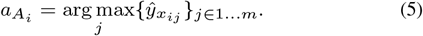

All of the following evaluation metrics are reported with the bag label *yxi*, the predicted bag label 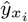 and the assigned allele *a_A_i*. When evaluating the performance per allele, we exclude the alleles with less than 50 either positive or negative samples.

## 3 Results

### 3.1 Self-supervised pretraining improves the prediction of MHC peptide interaction

Self-supervised pretraining has been shown to boost model performance for natural language processing and computer vision tasks (Devlin *et al.*, 2019; Chen *et al.*, 2020). Recent research has also shown the potential benefits of self-supervised pretraining on protein related tasks (Rao *et al.*, 2019; Nambiar *et al.*, 2020), such as contact prediction. However, to date, no work has explored self-supervised pretraining on MHC–peptide related tasks.

We first asked whether self-supervised pretraining helps with our prediction tasks. We investigated this in depth with the MHC–peptide binding affinity prediction task since all of the binding affinity values come from actual experiments, compared to presentation which entails sampling negatives. For this comparison, we binarized affinity values using the widely-used threshold of 500nM. We compared four strategies of training the same model architecture with the same hyperparameters:

- *Random*. Randomly initialized model trained end-to-end
- *Pretrain*. Model initialized with pretrained parameters from TAPE and trained end-to-end
- *Feature*. Model initialized with pretrained parameters from TAPE, but only the MLP classifier was trained. The BERT component was frozen during training and can be thought of as a kind of feature extractor.
- *Random Feature*. Randomly initialized model, but only the MLP classifier was trained

When comparing *Pretrain* with *Random*, the pretrained model not only achieved a better performance (auROC 0.872 vs. 0.853), but also converged much faster (Figure 2). Indeed, the pretrained model performed similarly as the best randomly initialized model after only a single training epoch (auROC 0.851 vs. 0.853). The pretrained model reached a good performance (auROC > 0.85) even at the first epoch whereas the randomly initialized model had poor performance at the start of the training process.

**Figure 2.**
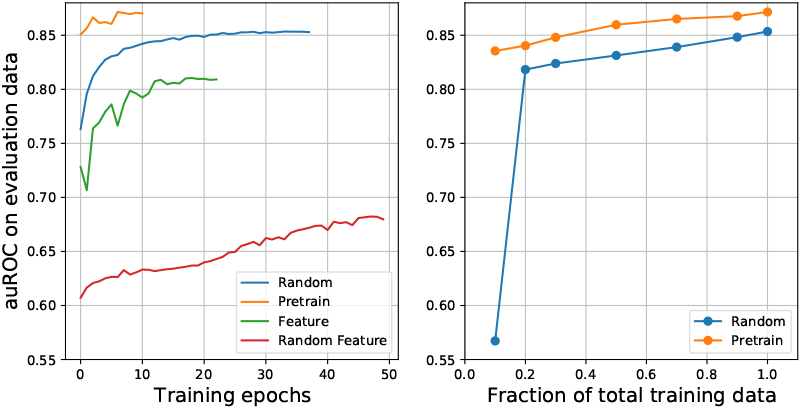
Comparing model performance with and without pretraining.

Based on these results, we use *Pretrain* for the remaining analysis.

### 3.2 Improved MHC class II binding prediction with transformers

We trained our MHC class II binding model with the dataset curated by Jensen *et al*. We used the same cross-validation folds so that peptides appearing in both training and testing folds are minimized. We compare our method (BERTMHC) against the sate-of-the-art class II peptide binding model NetMHCIIpan3.2 (Jensen *et al.*, 2018), which was trained with the same cross-validation splits. Our model outperformed NetMHCIIpan3.2 in terms of auROC for most of the alleles (48 out of 61, Figure 3). We also compared our model with PUFFIN (Zeng and Gifford, 2019b), which is a convolutional neural network model trained on the same dataset. BERTMHC (auROC = 0.8822) outperformed both PUFFIN-mean (auROC = 0.8774) and PUFFIN-BL (auROC = 0.8795) (Zeng and Gifford, 2019b).

**Figure 3.**
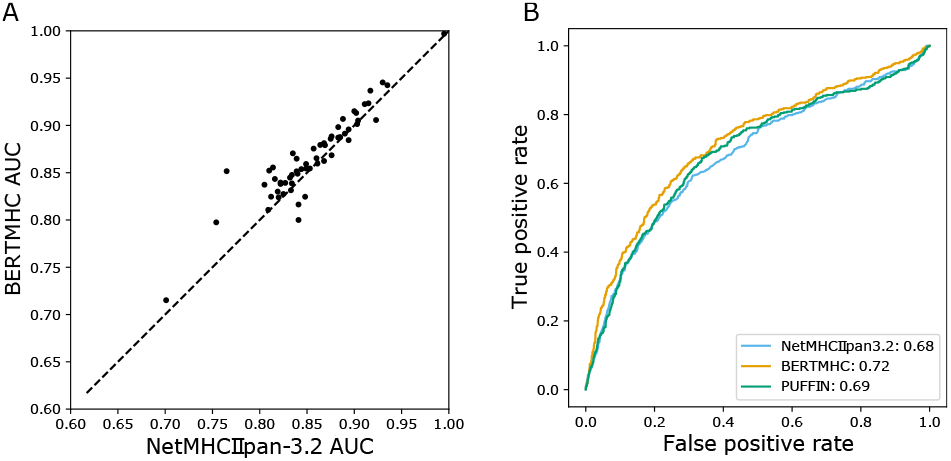
Comparing BERT with NetMHCIIpan3.2 under cross-validation. Each dot represents one MHC class II allele. The area under the receiver operating characteristic curve (auROC) for the BERT model (x-axis) are compared with the NetMHCIIpan3.2 model (y-axis).

We also used our independent MHC class II benchmark binding set, described in Section 2, to further benchmark the models. When evaluated on this independent data, our model (auROC = 0.72) performed better than NetMHCIIpan3.2 (auROC = 0.68) and PUFFIN (auROC = 0.69) in terms of classifying peptides into binders and non-binders.

### 3.3 Improved MHC class II presentation prediction with transformers

We then asked whether the same approach can be applied to train a better model to predict peptide presentation. Mass spectrometry is typically used to detect presentation of peptides on the cell surface. We trained our model on the curated mass spectrometry eluted data from Reynisson *et al.* (2020b), using the same cross-validation splits. We compared our model with the latest NetMHCIIpan4.0 model which was trained on the same dataset. Since the out-of-fold predictions of NetMHCIIpan4.0 were not provided, we compared our out-of-fold predictions against the prediction of the public version of NetMHCIIpan4.0, which is an ensemble of models trained on all cross-validation folds. Note that this comparison is in favor of the competing model since its training used our evaluation data.

For the MA data, we take the maximum prediction among the alleles in each bag of the prediction for that bag (see Methods). Evaluated on both SA and MA data, our model outperforms NetMHCIIpan4.0 in terms of both auROC and average precision (Table 1). Both models outperform NetMHCIIpan3.2; this is not surprising, since NetMHCIIpan3.2 is only trained on binding data (Table 1).

**Table 1.** Presentation performance on both MA and SA data, with all alleles combined

| Method | $\text{auROC}$ | Average Precision |
| --- | --- | --- |
| BERT | **$0.960 \pm 0.0004$** | **$0.835 \pm 0.001$** |
| NetMHCIIpan4.0 | $0.956 \pm 0.0004$ | $0.810 \pm 0.001$ |
| NetMHCIIpan3.2 | $0.704 \pm 0.001$ | $0.257 \pm 0.002$ |

We next evaluated the models on SA data only, where the labels are unambiguous. We observe that all models performed better in this setting compared with including the MA data in the evaluation (Table 2). Our model is consistently better than other models in terms of both auROC and average precision (Table 2). These results show that our model is able to perform well both when the allele labels are unambiguous and also when deconvolution is needed.

**Table 2.** Presentation performance on only SA data, with all alleles combined

| Method | $\text{auROC}$ | Average Precision |
| --- | --- | --- |
| BERT | **$0.972 \pm 0.0007$** | **$0.893 \pm 0.002$** |
| NetMHCIIpan4.0 | $0.961 \pm 0.002$ | $0.865 \pm 0.004$ |
| NetMHCIIpan3.2 | $0.796 \pm 0.001$ | $0.385 \pm 0.0025$ |

To further investigate the potential pros and cons of our model compared with others, we investigated the results for each allele. Our model outperformed NetMHCIIpan4.0 for more alleles than the other way around for both auROC (53 out of 86) and average precision (57 out of 89) when evaluated on both MA and SA data (Figure 4A). When evaluated on SA data only, we outperform NetMHCIIpan4.0 for most alleles in terms of both auROC (16 out of 19) and average precision (12 out of 19) (Figure 4B).

**Figure 4.**
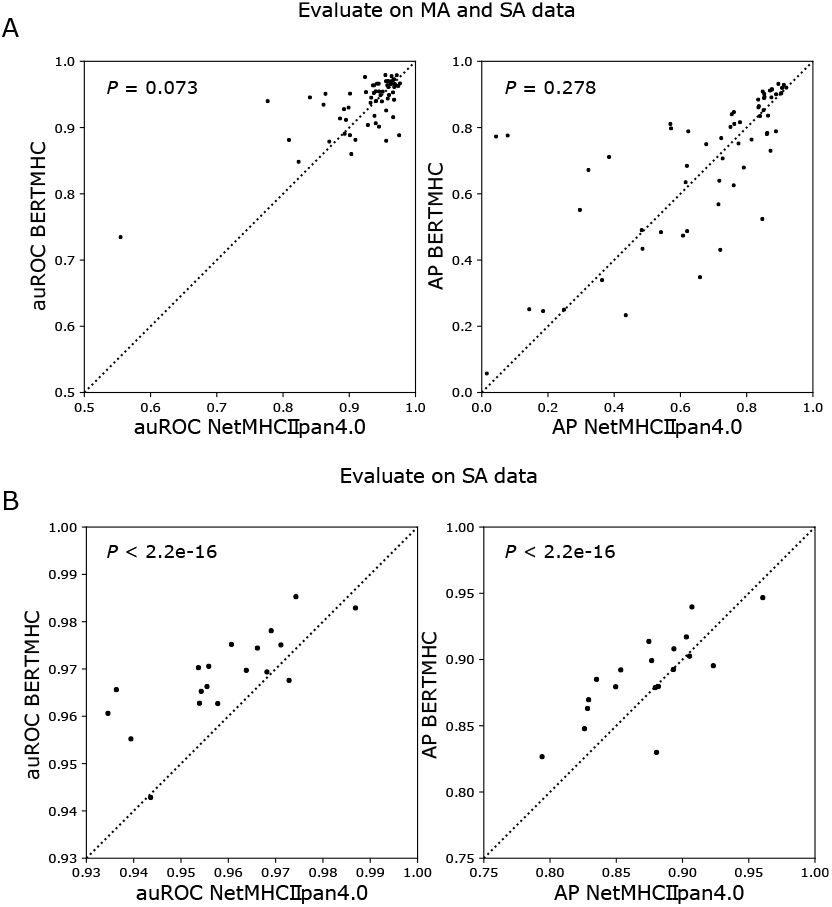
Comparing BERT with NetMHCIIpan4.0 under cross-validation. Each dot represents one MHC class II allele. (A) The auROC for the BERT model (x-axis) are compared with the NetMHCIIpan4.0 model (y-axis).

We next compared the methods on an independent mass spectrometry presentation dataset with no overlapping data from the training and evaluation set (see Methods). Our model outperforms NetMHCIIpan4.0 in terms of both auROC (0.89 versus 0.83) and average precision (0.60 versus 0.53) (Figure 5).

**Figure 5.**
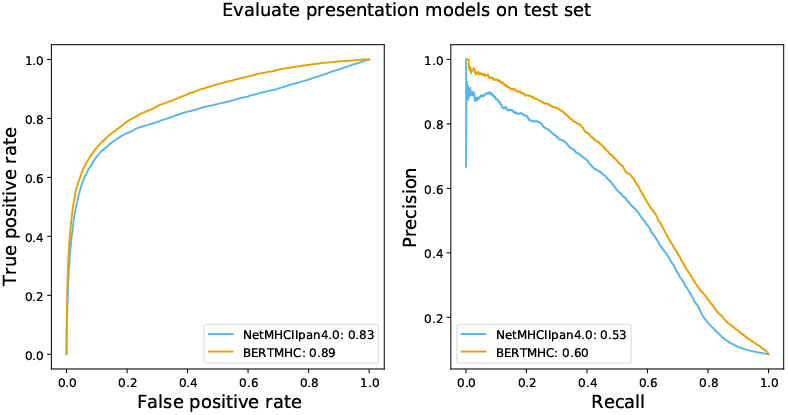
Comparing BERTMHC with NetMHCIIpan4.0 on independent mass spectrometry data from IEDB.

**Figure 6.**
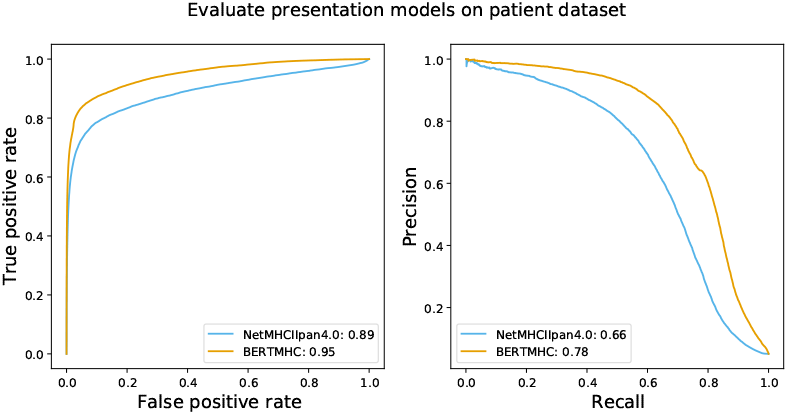
Comparing BERTMHC with NetMHCIIpan4.0 on mass spectrometry eluted peptides from 6 patient tumor samples.

To further evaluate our model under multiple instance setting, which is often the case when applying our model to real patients. We generated mass spectrometry data from peptides eluted from 6 patient samples (Methods). In total, 15, 277 unique peptides presented by HLA-DR molecules were eluted from 6 patients, with a median of 39, 64 peptide per patient. Each patient had two HLA-DR allele determined. We compared BERTMHC against NetMHCIIpan4.0 on this MA data. Negative peptides were randomly sampled from human proteome by matching the length distribution of positive peptides. We sampled 10 negative peptides for each positive peptide. Peptide scores were predicted as described in Equation 4 for both models. When evaluate on all patients combined, BERTMHC outperforms NetMHCIIpan4.0 both in terms of auROC (0.95 vs 0.89) and average precision (0.78 vs 0.66). When evaluated per patient, BERTMHC outperforms again NetMHCIIpan4.0 in all patients with maximum auROC improvement by 14.8% (Supplement Figure 1) and maximum average precision improvement by 61.0% (Supplement Figure 2).

## 4 Discussion

Predicting MHC–peptide binding has been a long standing problem, and many other approaches have also been developed. For example, embedding the input sequences for MHC-related tasks with self-supervised learning has also been explored in other directions. DeepLigand (Zeng and Gifford, 2019a) trained a language model from positive mass spectrometry peptides using the ELMo architecture (Peters *et al.*, 2018). The peptide embedding was shown to provide additional predictive information other than binding assays provide. Here we used a pre-trained network with a more effective architecture and trained from large corpus of protein sequences instead of only positive peptides. Moreover, the pretrained embedding model was applied to a concatenated sequence from both MHC and peptides, so that the model can potentially capture long distance interactions between MHC and peptide sequences.

Models with attention mechanisms have also been applied to MHC-peptide interaction prediciton tasks. ACME (Hu *et al.*, 2019) combines convolution and attention for MHC class I, MHCAttnNet (Venkatesh *et al.*, 2020) uses attention on top of a bidirectional LSTM (Venkatesh *et al.*, 2020) for both class I and class II. These models used one layer attention, for which the attention weights are more interpretable. Here we used multiple layers of self-attention, which is less interpretable but performed better in these tasks.

Despite the extensive existing work, previous models still have limited performance on MHC class II molecules. In this work, we have demonstrated that self-supervised pretraining for a transformer model leads to state-of-the-art performance for MHC class II binding and presentation prediction. We further provided a novel multiple instance learning strategy to address limitations of typical mass spectrometry assays for assessing eluted peptides. We conducted a thorough set of empirical experiments of the performance of the models for both binding and presentation. For presentation, we consider both the single allele setting, in which the exact MHC molecule to which a peptide is bound is known, as well as the multiple allele setting, in which only a set of possible MHCs to which a peptide was bound are known. In both cases, we show that our approach leads to state-of-the-art performance.

Our approach to predicting peptide presentation is solely based on the peptide and MHC sequences. Other studies have shown that the original context of the peptides and the peptide expression level are both important features for peptide presentation prediction (Reynisson *et al.*, 2020b; Chen *et al.*, 2019). Future work could integrate BERTMHC with gene expression to have more precise epitope prediction.

The proposed approach is applicable to other sequence-based predictions as well. We anticipate this approach to be useful to the community not only as a strategy for training binding and presentation models, but also as an approach to train protein sequence-based models for other challenges immunology, such predicting T-cell response. Considering that immunology assays are typically costly, a modeling strategy that improves data efficiency by performing self-supervised learning is valuable. The self-supervised pretraining step was not specific to downstream tasks, and was trained from generic protein sequences that may have very distinct biochemical properties compared to the sequences of our task. Nevertheless, the pretraining step was very beneficial. Preliminary work has been done to interpret the models trained from protein sequences with self-supervised learning (Heinzinger *et al.*, 2019; Vig *et al.*, 2020). This is a promising direction for future research in order to better understand what the models have learned and how that can guide better treatment decisions.

## Acknowledgements

We thank Carolin Lawrence, Timo Sztyler and Martin Renqiang Min for the discussions.

