## Supplementary Methods; Supplementary Table 1 for "BERTMHC: Improves MHC-peptide class II interaction prediction with transformer and multiple instance learning"

### 1 Supplementary Methods

#### 1.1 Data creation for patient mass spectrometry

We collected tumoral samples from **six** patients diagnosed with Non-Small Cell Lung Cancer (NSCLC) who were eligible for surgical resection. Tumor samples were rapidly snap frozen on liquid nitrogen upon collection. The tissue samples were lysed according to standard procedure with an Ultra Turrax (IKA Werke). The amount of tumor tissue was between 105 and 365 mg. From each of the tumor tissue lysates, HLA DR complexes were purified and analyzed by high resolution data-dependent acquisition mass spectrometry on a Q Exactive HF-X.

In order to identify peptides bound to the HLA DR molecules, resulting data was searched with MaxQuant against a database of human protein sequences from the reference proteome downloaded 26th June 2020 from UniProt (Consortium, 2019) website and containing 75 004 protein sequences.

The Class-II HLA types of each patient was determined by `seq2hla` tool (Boegel *et al.*, 2013) from RNA-seq data from tumor samples.

#### 2 Supplementary Tables

Table 1: Number of peptides per patient

| | Patient1 | Patient2 | Patient3 | Patient4 | Patient5 | Patient6 |
|---|---|---|---|---|---|---|
| # of peptides | 8,717 | 6,045 | 7,217 | 4,545 | 4,851 | 2,511 |

#### 3 Supplementary Figures

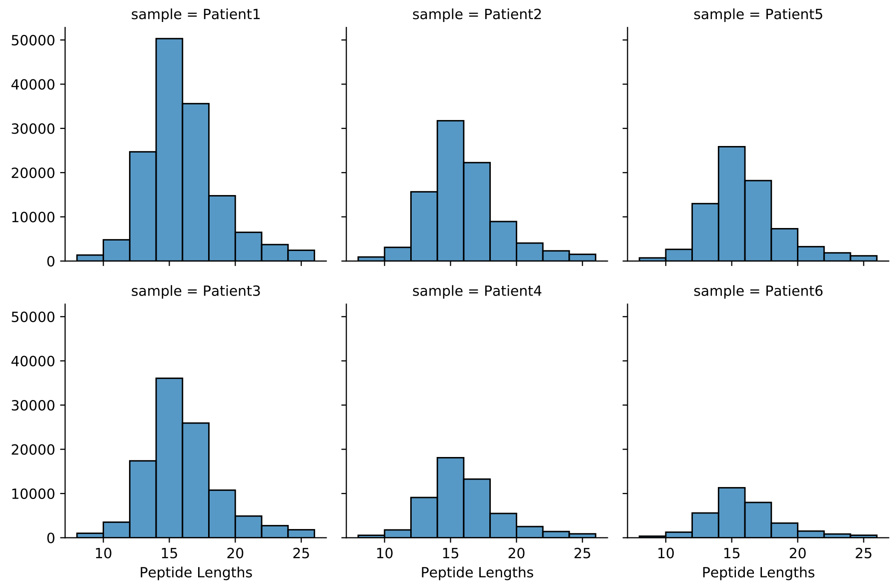


**Supplementary Figure** S1: Peptide length distribution for 6 patient samples.

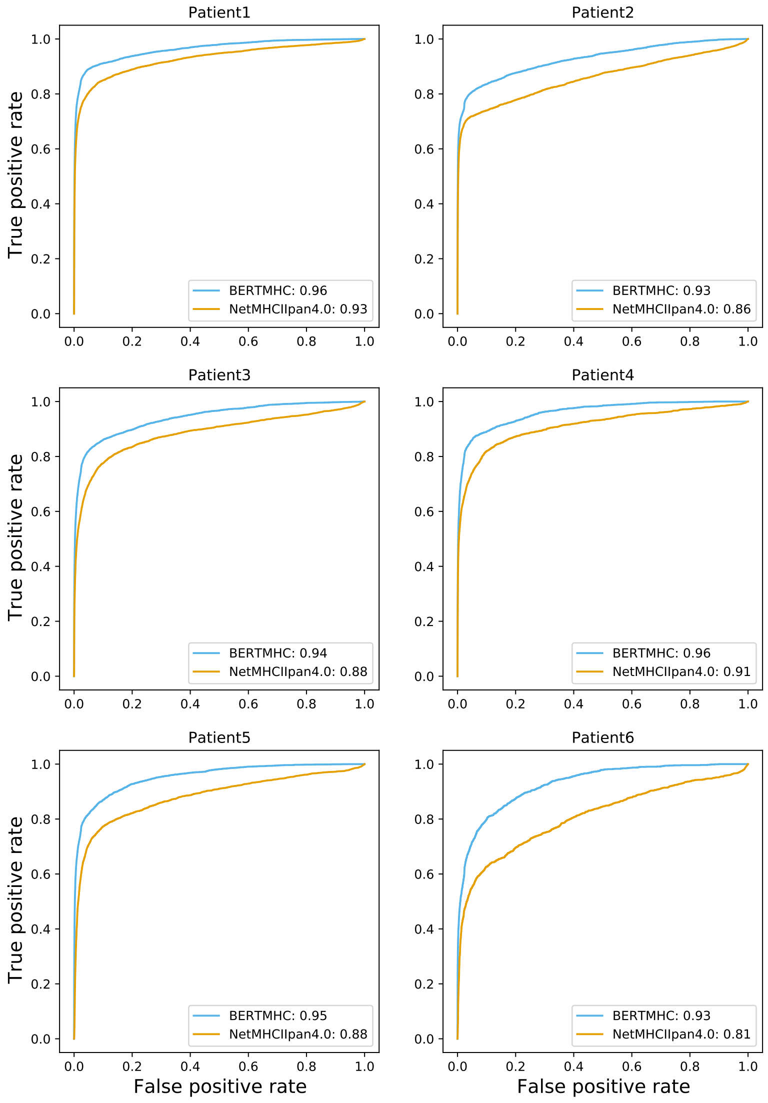


**Supplementary Figure** S2: Comparing BERTMHC with NetMHCIIpan4.0 on patient mass spectrometry data. Receiver operating characteristic curve plotted for BERTMHC (cyan) and NetMHCIIpan4.0 (yellow)

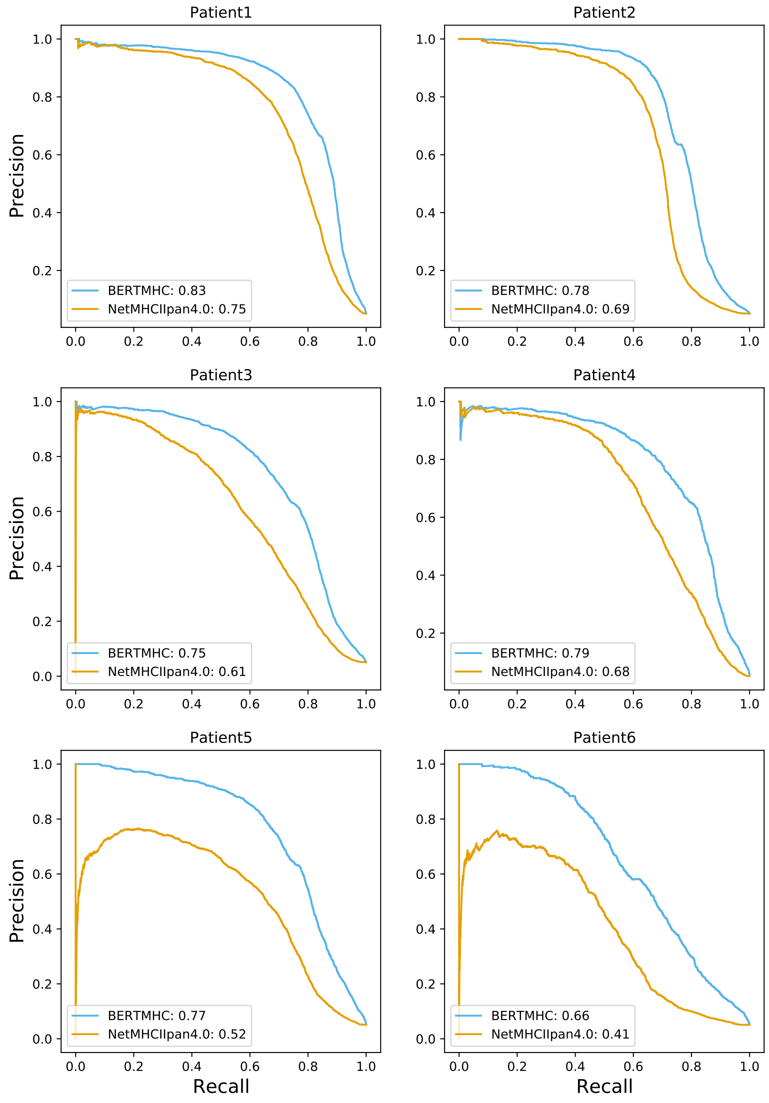


**Supplementary Figure** S3: Comparing BERTMHC with NetMHCIIpan4.0 on patient mass spectrometry data. Precision-recall curve plotted for BERTMHC (cyan) and NetMHCIIpan4.0 (yellow)
